# Myeloid Cell States in Influenza-Associated Pulmonary Aspergillosis Are Shaped by Iron Overload and Metabolic Reprogramming

**DOI:** 10.64898/2026.04.22.720189

**Authors:** Madeleine S. Grau, Brian P. Jackson, Carol Ringelberg, Fred W. Kolling, George S. Deepe, Robert A. Cramer, Joshua J. Obar

## Abstract

Virus-associated pulmonary aspergillosis is a life-threatening secondary infection that substantially increases morbidity and mortality in critically ill patients with respiratory virus infections. Influenza A virus (IAV) and SARS-CoV2 are known to disrupt pulmonary homeostasis, the mechanisms by which these perturbations render the host susceptibility to *Aspergillus fumigatus (Af)* remain incompletely understood. Here, we integrate an established murine model of influenza-associated pulmonary aspergillosis (IAPA) with single-cell RNA sequencing (scRNA-seq) to define the myeloid cell dysfunction that underlies IAPA establishment and progression.

Single-cell transcriptomic profiling of pulmonary monocytes and macrophages revealed that IAV-*Af* coinfection drives a marked shift away from interferon-mediated antiviral and antigen presentation programs toward stress-associated and redox-regulatory transcriptional states. Pathway analyses demonstrated coordinated suppression of phagocytic and interferon signaling pathways alongside enrichment of oxidative stress and mitochondrial metabolic signatures – changes that closely recapitulate transcriptional defects previously reported in human IAPA patients. Myeloid cells from IAV-*Af* coinfected mice further exhibited increased oxidative phosphorylation alongside reduced glycolytic and phagocytic activity, consistent with impaired antifungal effector function.

To elucidate how prior IAV infection generates a pulmonary microenvironment permissive to *Af* growth, we evaluated airway iron availability – a critical determinant of both fungal pathogenicity and immune regulation. IAV infection alone produced a significant elevation in bronchoalveolar iron levels accompanied by induction of iron-associated inflammatory mediators. Paradoxically, during IAV-*Af* coinfection, myeloid cells displayed markedly reduced expression of iron-sequestering and storage genes, revealing a fundamental disconnect between iron burden and cellular iron-handling capacity. Functionally, elevated iron accelerated *Af* germination and impaired macrophage-mediated fungal killing. Collectively, these findings identify IAV-induced pulmonary iron accumulation as a key driver of immunometabolic reprogramming in myeloid cells, resulting in compromised antifungal immunity and heightened susceptibility to secondary *Af* infection.

## Introduction

Severe respiratory virus infections, including those caused by influenza A virus (IAV) and SARS-CoV2, profoundly disrupt pulmonary homeostasis and predispose patients to a broad range of secondary infections [1–6]. Although these viruses were originally recognized primarily as drivers of susceptibility to secondary bacterial superinfections, secondary fungal infections have emerged as an increasingly serious and often underappreciated threat [7–11]. The ubiquitous environmental mold *Aspergillus fumigatus (Af),* a well-established cause of invasive pulmonary aspergillosis in severely immunocompromised patients, is now recognized as a significant complication among ICU-admitted patients with severe influenza or COVID-19 [6, 8, 11–14]. Influenza-associated pulmonary aspergillosis (IAPA) carries mortality rates of 30-60% – roughly double that of critically ill influenza patients without aspergillosis [8, 10, 15–17] – underscoring the urgent need to understand the pathogenic mechanisms that drive this coinfection syndrome.

The mechanisms by which IAV disrupts pulmonary defenses to enable bacterial superinfections are well characterized and include epithelial barrier disruption, impaired mucociliary clearance and dysregulated innate immune cell responses [5, 18–21]. More recently, several clinically relevant murine models of IAPA have been developed, enabling systematic investigation of the host-pathogen interactions that govern susceptibility to secondary fungal infection [22–27]. Collectively, these studies have identified at least three interconnected mechanisms by which prior IAV infection facilitates *Af* establishment: (1) viral-induced epithelial disruption that compromises physical barrier defenses [25, 27]; (2) impaired antifungal phagocytic and killing functions of myeloid cells [22]; and (3) defective neutrophil-mediated extracellular killing mechanisms [23, 26]. Severe inflammatory responses and cytokine storm have also been described in IAPA patients [28] and are recapitulated in murine models. Our group previously demonstrated that myeloid cells exhibit reduced antifungal activity during IAPA, with defective *Af* phagocytosis in neutrophils and monocytes leading to reduced fungal clearance [22] – findings that align with clinical observational studies of innate immune populations in human IAPA [29–31]. Nevertheless, a comprehensive mechanistic understanding of myeloid cell dysfunction following IAV infection remains elusive.

Competition for iron between host and pathogen is a fundamental determinant of microbial virulence and disease outcome [32, 33]. Under homeostatic conditions, the host restricts free iron availability through sequestration mechanisms that collectively limit pathogen access to this essential micronutrient – a strategy termed nutritional immunity. Viral infections are known to perturb iron homeostasis in ways that persist well beyond the acute phase of infection, with significant consequences for host susceptibility to secondary pathogens. Iron-overloaded macrophages arising in the context of viral infection have been shown to impair pathogen resolution and promote immune dysfunction [34]. *Af,* like other pathogens, requires iron as an essential cofactor for the fundamental biochemical processes [35, 36], and critically, *Af* has evolved sophisticated mechanisms to adapt to and exploit ambient iron concentrations through indirect uptake pathways [37]. Despite growing evidence that viral infection induces pulmonary iron overload, the potential link between IAV-driven iron dysregulations and susceptibility to secondary *Af* infection has not been directly investigated.

In the present study, we address this gap by integrating our established murine IAPA model [22] with high-resolution single-cell RNA sequencing (scRNA-seq) to define the myeloid cell transcriptional landscape across infection conditions. Through this approach, we validate that our murine model faithfully recapitulates the conserved myeloid dysfunction signatures associated with impaired antiviral and antifungal responses observed in human IAPA and COVID-19-associated pulmonary aspergillosis (CAPA) datasets. Importantly, we demonstrate that IAV infection drives significant pulmonary iron accumulation that coincides with defective myeloid cell iron handling and impaired fungal clearance following secondary *Af* challenge. Mechanistically, elevated airway iron promotes *Af* germination and directly impairs myeloid cell antifungal function. Together, these findings position iron dysregulation as a novel, potentially targetable disease-initiating factor in IAPA, with broad implications for host-directed therapeutic strategies in severe respiratory virus infections.

## Results

### Single-cell RNA sequencing characterizes dynamic myeloid cell remodeling across pulmonary infection states

We previously demonstrated that pulmonary macrophages exhibit impaired antifungal responses following IAV infection, manifesting as reduced phagocytic killing and diminished *Af* spore uptake [22]; however, the transcriptional programs driving this dysfunction had not been systematically characterized. To address this comprehensively, we performed single-cell RNA sequencing of monocyte and macrophage populations isolated from the lungs of mice subjected to four experimental conditions: naïve (PBS/PBS), IAV only (IAV/PBS), *Af* only (PBS/*Af*), IAPA coinfection (IAV/*Af*). Mice were intranasally inoculated with IAV strain A/Puerto Rico/8/34 (100 EID₅₀) on day 0, followed by intratracheal challenge with 3.5 × 10⁷ *A. fumigatus* CEA10 conidia at day 6, and lungs were harvested 24 hours after secondary challenge.

To capture the transcriptional heterogeneity of myeloid populations within the infected lung microenvironment, we specifically isolated monocytes and interstitial macrophages while excluding alveolar macrophages -- a decision informed by prior findings demonstrating that alveolar macrophages do not exhibit significant changes in antifungal killing capacity during IAPA coinfection [22], and that their abundance could bias downstream analyses (Supplemental Figure 1A). Cell isolation employed CD11b-positive magnetic enrichment followed by a two-step FACS strategy to exclude non-target populations based on lineage markers [38]. Lung tissue-derived CD45^(⁺)^ immune cells were initially positively selected using CD45 labeling. Subsequently, a negative depletion strategy was applied using intravenous CD45 (to exclude circulating cells), Ly6G (neutrophils), NK1.1 (NK cells), CD4 and CD8 (T cells), CD19 (B cells), and Siglec-F (alveolar macrophages). The remaining cell population was then positively sorted for both CD64 and CD11b expression to enrich for tissue-resident macrophages and recruited monocytes (Supplemental Figure 1B). A second FACS sort was performed after DAPI spiking to select live cells ensuring high viability for downstream sequencing (Supplemental Figure 1C). Approximately 10,000 viable DAPI^(-)^ CD11b^(+)^ CD64^(+)^ cells from five pooled mice per condition were sequenced using the 10x Genomics Chromium platform, and single-cell transcriptomes from all four conditions were integrated for cross-condition comparison.

Unbiased clustering of the integrated dataset identified 11 transcriptionally distinct clusters representing diverse myeloid activation states and infection-specific responses (Figure 1A). Expression of canonical lineage-defining monocyte and macrophage markers was confirmed across all clusters (Figure 1B), and negatively selected features were absent from sequenced cells (Supplemental Figure 2), validating the specificity of our isolation strategy. Sequenced CD11b^(+)^ CD64^(+)^ cells were largely negative for *Ly6g, Klrb1c* (NK1.1)*, Cd4, Cd8a, Cd19,* and *Siglecf* indicating successful isolation of the pulmonary myeloid population. All clusters also exhibited detectable levels of *Fcgr1* (CD64) and *Itgam* (CD11b), consistent with their positive selection during the isolation process (Figure 1B). High expression of *Ly6c* and *Plac8* across most clusters supported their classification as inflammatory monocytes or macrophages, while variable expression of *Cd14*, *Fcgr3* (CD16), and *Irf8* across clusters indicated a continuum of activation states and differentiation trajectories (Figure 1B).

**Figure 1.**
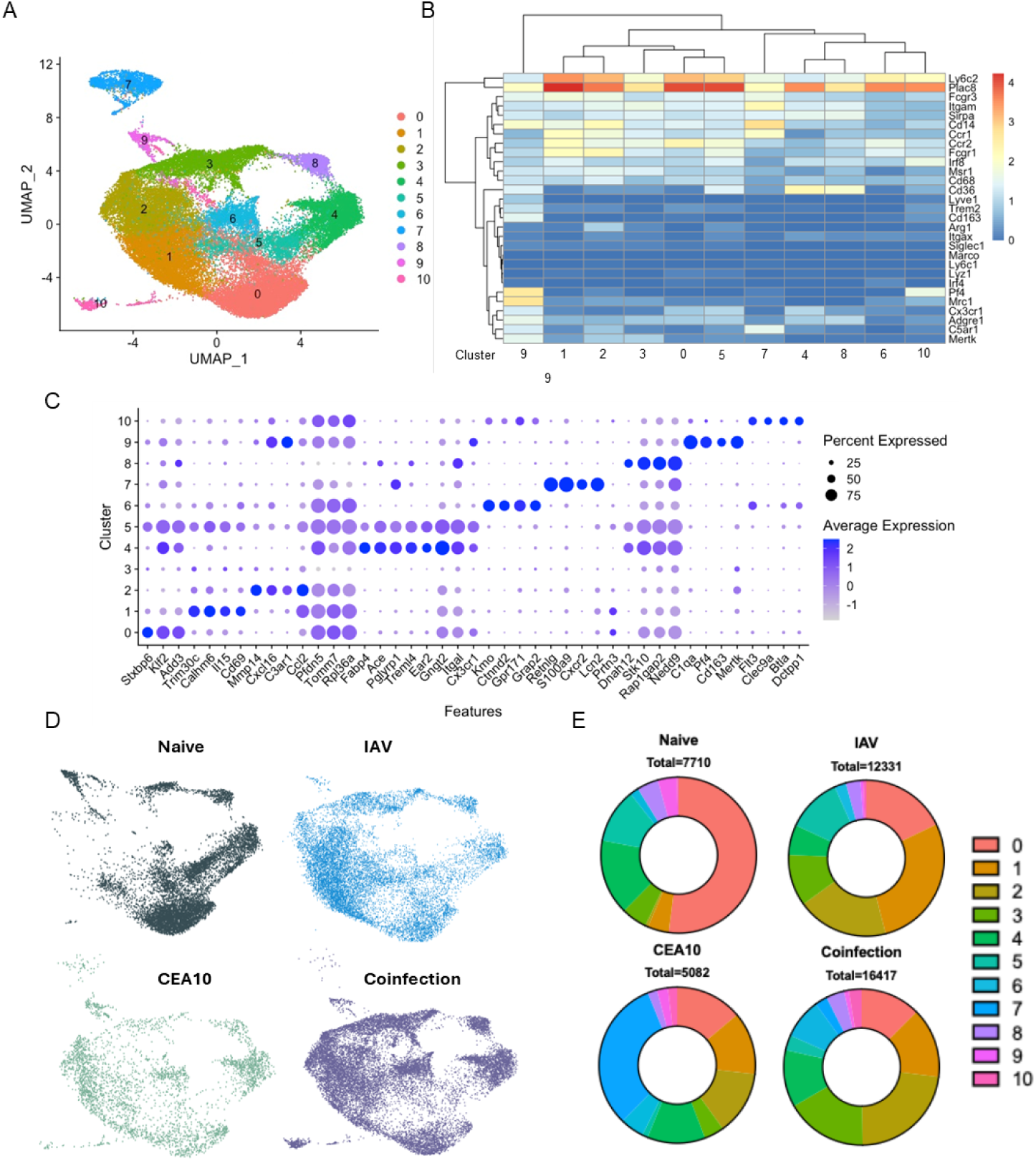
Single-Cell transcriptional landscape of lung monocytes/macrophages in IAPA model. (A) Uniform Manifold Approximation and Projection (UMAP) visualization of integrated single-cell RNA sequencing data generated using the 10x Genomics platform, showing 11 distinct myeloid cell clusters identified from lung monocyte and macrophage populations. (B) Heatmap showing logarithmic expression of canonical monocyte/macrophage lineage and activation marker genes across identified clusters. Gene expression values are scaled across clusters for visualization. (C) Dot plot showing expression levels of selected marker genes across each cell cluster. Dot size represents the proportion of cells expressing the gene within the cluster and color intensity represents the scaled average expression level. (D) Condition-split UMAPs showing the distribution of cells originating from each experimental condition: Naïve, IAV, CEA10, and Coinfection. (E) Donut plots showing the relative proportions of myeloid cell clusters across experimental conditions (Naïve, IAV, CEA10, and Coinfection). The total number of cells contributing to each condition following quality control filtering and dataset integration is indicated above each plot.

Differential gene expression (DEG) analysis revealed distinct transcriptional programs that enabled functional grouping of clusters (Figure 1C). The dataset comprised a heterogeneous mixture of inflammatory monocytes, tissue-resident interstitial macrophages, neutrophil-like myeloid cells, and DC-like populations. Cluster 0, predominantly composed of cells from naïve lungs, expressed genes associated with cellular homeostasis, consistent with a resting myeloid state. Clusters 1, 2, 4, 5, 7 and 9 displayed elevated expression of activation and inflammatory markers including *Ccl2, Pglyrp1, Mmp14, Il15, Cd69,* and *C3ar1*, indicative of robust pro-inflammatory activation. Clusters 4, 5, and 9 co-expressed *C1qa, Cd163, Mertk, Pf4,* and *Rap1gap2*, markers characteristic of tissue-resident myeloid identity. Clusters 6 and 8 were enriched for genes involved in immune signaling and regulation. Clusters 7 and 10 exhibited highly distinctive transcriptional programs: Cluster 7 expressed canonical neutrophil-associated genes including *Retnlga, S100a8, S100a9, Lcn2, Ngp, Cxcr2, Cxcl2,* and *Cxcl3*, consistent with a granulocytic or neutrophil-like transcriptional state, while Cluster 10 expressed dendritic cell–associated markers *Cd24a* and *Clec9a*, consistent with conventional DC populations. These findings underscore the high degree of context-dependency in myeloid cell responses to pulmonary infection.

In addition to transcriptional profile differences in the myeloid cell clusters, UMAP projection of cells stratified by experimental condition revealed marked, condition-specific shifts in cluster distribution (Figure 1D). Cluster 0 predominated in naïve lungs, Cluster 1 in IAV-infected lungs, and Cluster 7 in *Af*-infected lungs. In contrast, cells from the IAPA coinfection condition were broadly distributed across Clusters 0, 1, 2, and 3, with no single cluster dominating – a pattern indicative of a heterogeneous, dysregulated transcriptional response (Figure 1E). Notably, the proportion of Cluster 7 cells was markedly reduced in the IAPA coinfection group relative to the *Af-*only condition. This condition-specific redistribution of myeloid states during IAPA coinfection suggests impaired coordination of antiviral and antifungal responses and provides the transcriptional foundation for subsequent mechanistic analyses.

### Murine IAPA transcriptome mirrors pathogenic) phenotypes observed in human IAPA patients

We next sought to contextualize transcriptional differences between our IAV infection and IAPA coinfection cohorts by comparing our murine scRNA-seq dataset to previously characterized immune defects in human and murine IAPA [30, 31]. Differential expression analysis between the IAV infection and IAPA coinfection groups revealed a marked and coordinated shift away from interferon-driven antiviral programming toward a transcriptional state dominated by tissue injury, stress adaptation and metal ion handling (Figure 2A). Genes preferentially expressed in the IAV condition were enriched for interferon-stimulated and antiviral response genes -- including *Cxcl9*, *Ifit1*, and *Rsad2* --consistent with a robust type I interferon–driven antiviral program. In contrast, genes upregulated during IAPA coinfection included markers of tissue injury and stress responses (*Ngp*, *Ccl17*, *Camp*, *S100a9*) as well as metal stress adaptation (*Calm4, Mmp9, S100a9, Mt3*, *Adam23*). These data indicate that secondary *Af* challenge fundamentally reshapes myeloid transcriptional landscape, redirecting cells away from antiviral immunity toward programs associated with tissue damage, inflammation, and impaired effector function – a pattern that aligns with immune dysregulation described in human IAPA.

**Figure 2.**
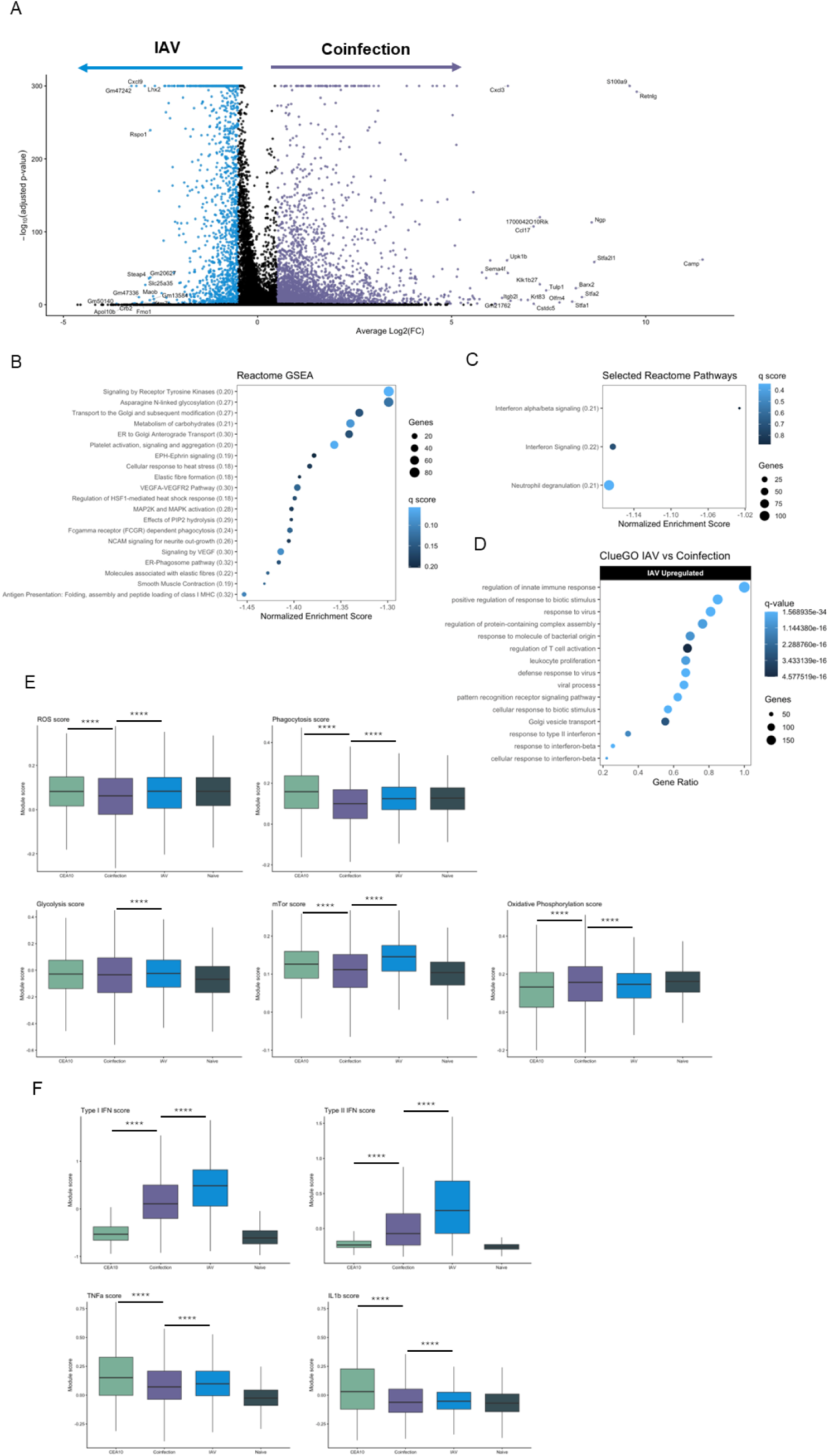
Coinfection suppresses antiviral programs and reprograms myeloid cell metabolism toward stress-associated state. (A) Volcano plot showing differential gene expressions comparing IAV and coinfected cells. Genes upregulated in IAV are shown on the left (blue) and genes upregulated in coinfection are shown on the right (purple). The top 20 significantly differentially expressed genes are annotated. (B) Dot plot showing the top 20 significant pathways identified by Reactome gene set enrichment analysis (GSEA) for genes upregulated in IAV compared with IAPA (BH-adjusted q-value < 0.25). (C) Reactome GSEA dot plot displaying selected pathways corresponding to those reported in Feys et al. (D) ClueGO pathway enrichment analysis of genes downregulated in coinfection relative to IAV. (E) Box plots showing module scores for curated functional gene sets across naïve, IAV, CEA10, and coinfection conditions, calculated using KEGG, Hallmark, or Reactome ontology gene sets (gene lists provided in the Appendix). (F) Box plots showing module scores for cytokine-responsive transcriptional pathways, including TNF-α, IL-1β, type I interferon (IFN-α/β), and type II interferon (IFN-γ), across experimental conditions. Adjusted *p*-values were calculated using the Wilcoxon rank-sum test.

To enable direct comparison with human transcriptional data, we performed Reactome pathway analysis guided by the framework of Feys *et al.* [30]. Unbiased analysis of the top 20 downregulated pathways in IAPA coinfection versus IAV infection revealed coordinated suppression of phagocytic programs, including antigen processing and presentation, the ER–phagosome pathway, and Fcγ receptor–mediated phagocytosis (Figure 2B). Although individual pathways did not uniformly reach stringent multiple-testing corrected significance thresholds, the coordinated directional enrichment across these related pathways strongly suggests attenuation of phagocytic and antigen-processing capacity during IAPA coinfection. When pathway results were filtered to those highlighted in the Feys *et al* human IAPA dataset [30], three of the six pathways reported in patients – neutrophil degranulation, type I interferon signaling, and interferon-γ signaling – showed concordant enrichment trends in our murine dataset (Figure 2C), providing cross-species validation of the model.

Complementary ClueGO analysis of the top 15 downregulated pathways in IAPA coinfection versus IAV infection further corroborated these findings, revealing enrichment of gene ontology terms related to antiviral defense (Figure 2D) – a signature that significantly overlaps with those reported from BAL fluid cells of human IAPA patients [31]. ClueGO also identified suppressed response to type I and type II interferons, paralleling interferon-pathway attenuation in human disease, alongside downregulation of innate immune response regulation, pattern recognition receptor signaling, and leukocyte proliferation. Together, Reactome and ClueGO analyses confirm a conserved and coordinated suppression of antiviral and interferon-driven transcriptional programs during IAPA coinfection.

To quantitatively validate these pathway-level changes across all conditions, we calculated module scores for curated pathway gene sets (Figure 2E). IAPA coinfection samples showed reduced scores for reactive oxygen species (ROS) production (0.0579 ± 0.0009 vs. IAV: 0.0711 ± 0.0010 and *Af*: 0.0800 ± 0.0014; p < 0.0001) and phagocytosis (0.0977 ± 0.0008 vs. IAV: 0.1270 ± 0.0008 and *Af*: 0.1580 ± 0.0016; p < 0.0001) relative to either single-infection condition, indicating broadly blunted innate effector function. Additionally, myeloid cells from IAPA coinfected lungs exhibited a characteristic metabolic shift: elevated oxidative phosphorylation scores (0.1460 ± 0.0010 vs. 0.1360 ± 0.0009; p < 0.0001) coupled with reduced mTOR signaling (0.1070 ± 0.0005 vs. 0.1370 ± 0.0005; p < 0.0001) and glycolysis scores (-0.0359 ± 0.0014 vs. -0.0224 ± 0.0015; p < 0.0001) relative to IAV infection alone. This metabolic configuration – favoring oxidative phosphorylation over mTOR-dependent glycolysis – is consistent with dysfunctional antifungal phenotype, as glycolytic reprogramming has been identified as a prerequisite for efficient fungal killing by myeloid cells [31].

Finally, to quantify downstream signaling of cytokines central to antifungal and antiviral immunity (IL-1β, TNF-α, type I IFN, and type II IFN), we generated cytokine-responsive pathway module scores (Figure 2F) and compared these to the patterns reported in the human IAPA cohort by Feys *et al.* [30]. Across all four cytokine pathways, IAPA coinfection samples exhibited significantly lower scores than IAV-only samples (IL-1β: - 0.0620 ± 0.0012 vs. -0.0531 ± 0.0011, p<0.01; TNF- α: 0.0984 ± 0.0015 vs. 0.1080 ± 0.0014, p < 0.0001; type II IFN: 0.0298 ± 0.0026 vs. 0.3580 ± 0.0047, p < 0.0001; type I IFN: 0.1660 ± 0.0036 vs. 0.4440 ± 0.0044, p < 0.0001), indicating a broad failure to mount or sustain cytokine-driven transcriptional programs during coinfection. The reduced IL-1β, and TNF-α module scores in IAPA coinfection relative to both IAV and *Af* single-infection conditions further support a model in which secondary fungal challenge dampens myeloid cell responsiveness to key proinflammatory cytokines, thereby constraining both antifungal and antiviral defenses. Taken together, these results demonstrate that our murine IAPA model faithfully recapitulates the hallmark transcriptional defects of human IAPA – namely, loss of interferon signaling, impaired phagocytic and cytokine-driven programs, and adoption of a stress-associated metabolic state.

### scRNA-seq analysis of myeloid cells from IAV-infected mice reveals enrichment of iron-related stress pathways

Because initiation of IAPA coinfection depends on the pulmonary microenvironment into which *Af* spores are deposited, we next examined how IAV infection reshapes monocyte and macrophage transcriptional states relative to the naïve lung. IAV infection is well established to damage airway epithelium and alter the available nutrient milieu – including glucose, lactate, and free metal concentrations – in ways that may facilitate fungal germination and outgrowth [39]. We therefore used our scRNA-seq dataset to characterize the transcriptional state of pulmonary myeloid cells in naïve and IAV-infected lungs, corresponding to the pulmonary conditions present at the time of *Af* conidial deposition in our model.

Gene set enrichment analysis (GSEA) confirmed that IAV infection robustly activated monocyte and macrophage antiviral programs, with the most strongly upregulated pathways involving innate immune activation, sensing of viral nucleic acids, intracellular signaling cascades, and cytokine-mediated communication (Figure 3A). These findings confirm that lung myeloid cells mount appropriate and competent antiviral responses during isolated IAV infection – establishing the immunological context into which *Af* conidia are subsequently introduced.

**Figure 3.**
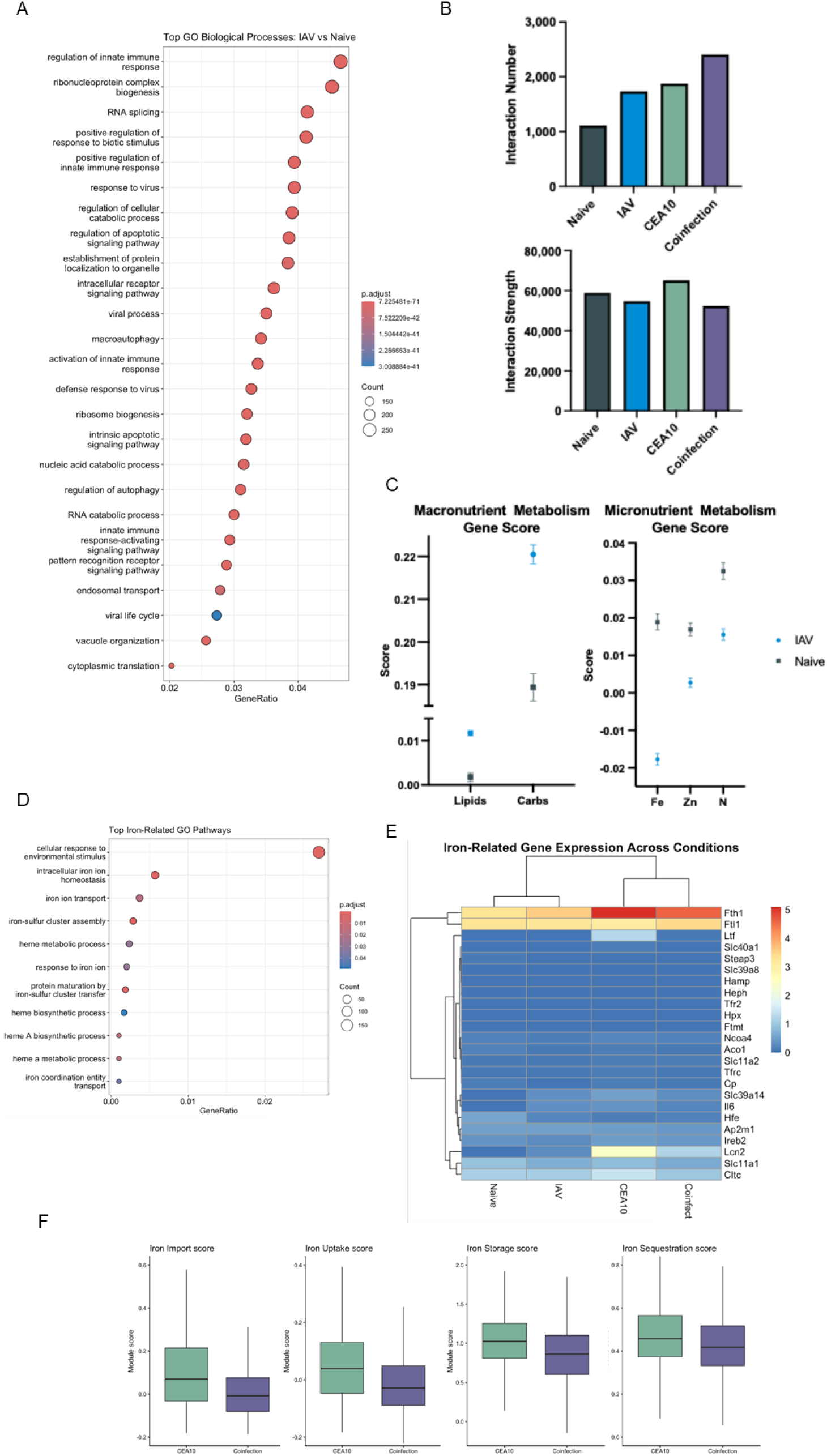
Influenza infection induces iron-responsive transcriptional pathways in pulmonary monocytes and macrophages. (A) Gene Ontology (GO) enrichment analysis showing the top 25 enriched biological processes identified from differential gene expression analysis comparing IAV-infected cells to naïve controls. (B) CellChat-based inference of intercellular communication across naïve, IAV, CEA10, and coinfection conditions. Bar plots depict the total number of inferred ligand–receptor interactions (top) and the aggregate interaction strength across signaling pathways (bottom). (C) Module scores calculated from curated gene sets representing macronutrient (lipid and carbohydrate) and micronutrient (iron, zinc, and nitrogen) metabolic processes. Data are shown as mean ± SEM. Gene lists used for module scoring are provided in the Appendix. (D) Curated Gene Ontology enrichment analysis of significantly enriched iron-associated biological processes identified from differential expression analysis comparing IAV and naïve controls. (E) Heatmap showing row-scaled expression of iron-related genes across experimental conditions (Naïve, IAV, CEA10, and Coinfection). Gene expression values were hierarchically clustered. (F) Box plots showing iron-related pathway module scores for CEA10 and coinfection cells. Adjusted *p*-values were calculated using the Wilcoxon rank-sum test.

To evaluate the broader landscape of cell-cell communication during IAPA, we applied CellChat [40] to infer ligand–receptor interaction networks across all four infection conditions (Figure 3B). Although IAPA coinfection yielded a greater number of inferred ligand-receptor interactions compared with either single-infection condition, the aggregate signaling strength across pathways was markedly reduced. This paradoxical pattern – increased communicative complexity alongside diminished signaling potency – suggests that IAPA coinfected myeloid cells retain the structural framework for communication but exhibit profoundly impaired signaling efficiency, consistent with functional exhaustion within the coinfection microenvironment.

Given the established capacity of IAV to drive epithelial injury and disrupt local metal homeostasis [39, 41, 42], and the critical importance of iron as a micronutrient for *Af* pathogenesis [35, 43, 44], we next specifically examined nutrient-associated transcriptional programs. Using curated gene sets, we evaluated pathway-level signatures associated with macronutrient (lipid and carbohydrate) and micronutrient (iron, zinc, and nitrogen) metabolism in naïve and IAV-infected lungs (Figure 3C). While carbohydrate metabolism scores increased during IAV infection (0.2210±0.0023 vs 0.1890±0.0032, p<0.0001), micronutrient-related transcriptional programs showed the opposite pattern. Gene signatures associated with iron (-.0.0117±0.0015 vs 0.0189±0.0021, p<0.0001), zinc (0.0027±0.0012 vs 0.0169±0.0039, p<0.0001), and nitrogen metabolism (0.0155±0.0015 vs 0.0325±0.0022, p<0.0001) were significantly reduced in IAV-infected cells compared to naïve controls, indicating dysregulated micronutrient handling during viral infection.

To further interrogate iron-specific responses, we preformed GO analysis comparing naïve and IAV-infected conditions, which reveal significant enrichment of pathways governing intracellular iron ion homeostasis and related iron-handling mechanisms (Figure 3D) – confirming that IAV infection alone induces a measurable iron-regulatory response in lung myeloid cells. Assessment of individual iron-related genes between naïve and IAV-infected samples were modest, IAPA coinfection was associated with pronounced reductions in key iron-regulatory genes relative to *Af* infection alone --including *Fth1* (ferritin heavy chain) and the iron-binding and sequestration factors *Ltf* and *Lcn2*. These reductions indicate that myeloid cells in the IAPA coinfected lung have substantially diminished capacity to store or neutralize excess iron.

To quantify these transcriptional changes at the pathway level, we performed gene set scoring for monocyte and macrophage iron-related functions encompassing iron uptake, import, storage, and sequestration (Figure 3F). IAPA coinfected samples exhibited significantly reduced scores in iron uptake (-0.0110±0.0008 vs 0.0507±0.0017, p<0.0001), import (0.0118±0.0009 vs 0.0992±0.023, p<0.0001), and storage (0.837±0.003 vs 1.020±0.005, p<0.0001) pathways compared to the *Af*-only group, with moderate reduction also observed in iron sequestration (0.420±0.001 vs 0.466±0.002, p<0.0001). These findings confirm that iron-regulatory responses are paradoxically impaired during IAPA coinfection despite the increased likelihood of iron overload within the damaged pulmonary microenvironment. The combination of elevated extracellular iron burden and a failure of cellular iron-handling responses – together with blunted myeloid intercellular communication -- likely contributes to early fungal outgrowth and the immune dysfunction that is characteristic of IAPA.

### IAV infected mice exhibit increased airway iron levels and induction of iron-regulatory mediators

To directly assess whether viral infection drives micronutrient accumulation in the lung, we measured total iron concentration in bronchoalveolar lavage fluid (BALF) collected from naïve or IAV-infected mice at day 6 post-challenge – the time point corresponding to secondary *Af* inoculation in our IAPA model. ICP-QQQ-MS analysis revealed significantly elevated iron concentrations in IAV-infected mice compared with uninfected controls (0.485 ± 0.102 µg/ml vs. 0.152 ± 0.018 µg/ml; p < 0.0001), confirming that IAV infection produces substantial pulmonary iron overload (Figure 4A). These findings are consistent with the established paradigm that viral-induced epithelial damage releases intracellular iron stores and broadly perturbs metal homeostasis. Additional biologically relevant metals (zinc, nitrogen, copper, and manganese) were also elevated to a lesser but statistically significant degree (Supplemental Figure 3), suggesting that IAV-associated injury broadly disrupts the nutritional landscape of the lower respiratory tract.

**Figure 4.**
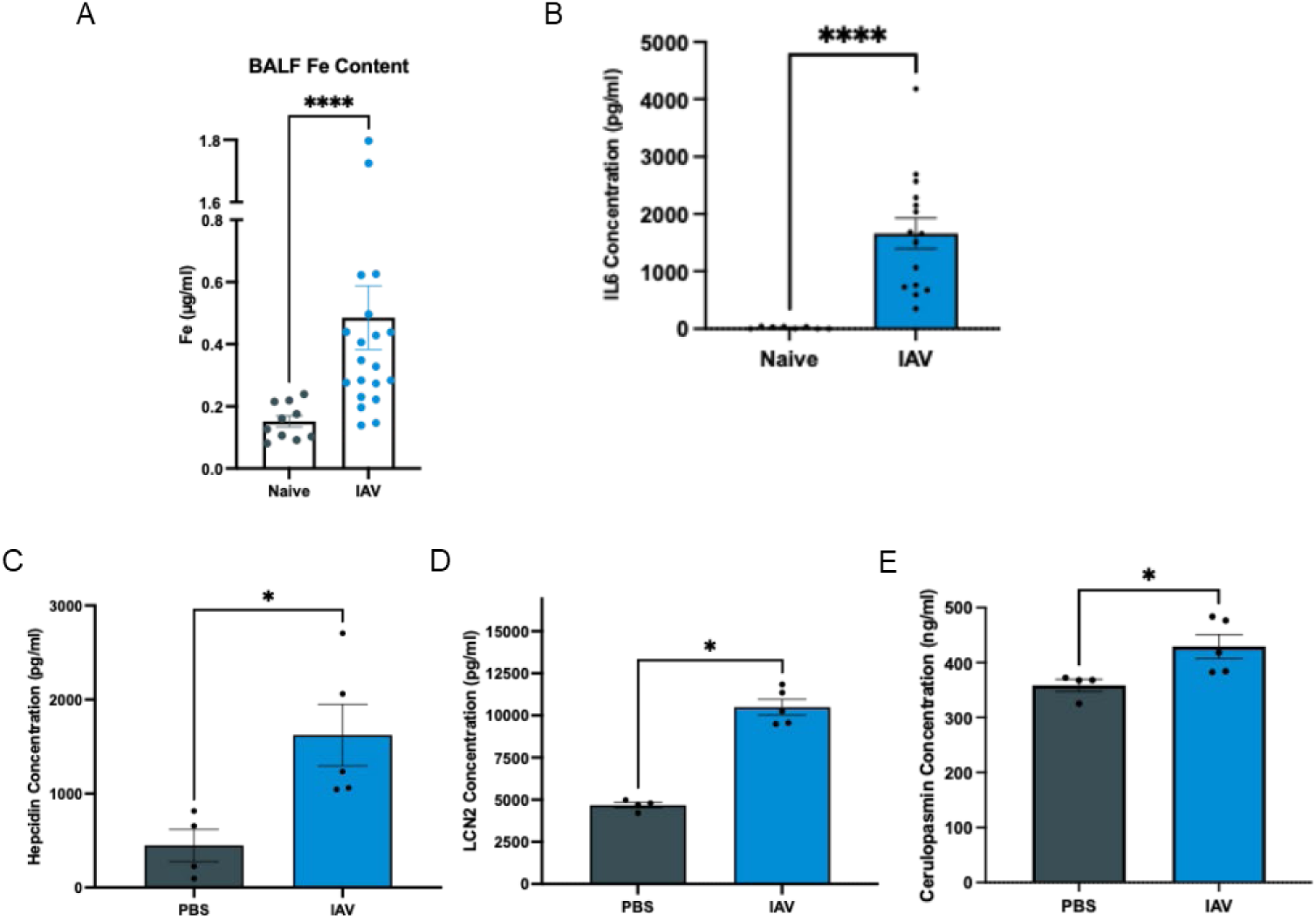
Influenza infection induces pulmonary iron accumulation. Total iron concentration and iron-regulatory mediators were measured in bronchoalveolar lavage fluid (BALF) collected from naïve (PBS-treated) and influenza A virus (IAV)-infected mice at day 6 post-infection. (A) Total iron concentration in BALF was quantified by ICP-QQQ-MS. Data were pooled from two independent experiments. (B) Interleukin-6 (IL-6) concentrations, (C) hepcidin, (D) Lipocalin-2 (LCN2) and (E) ceruloplasmin concentrations in BALF measured by ELISA. Bars represent mean ± SEM, with individual points indicating biological replicates. Statistical significance was determined using unpaired two-tailed Mann–Whitney U tests. p < 0.05 (*)

We next asked whether this iron accumulation was accompanied by induction of iron-responsive immune mediators. IAV infection led to robust upregulation of the iron-associated cytokine IL-6 (1662 ± 267 pg/ml vs 18.8 ± 6.9pg/ml; p < 0.0001) (Figure 4B), a well-established driver of acute-phase iron redistribution and hepcidin production [45, 46]. Correspondingly, the iron-regulatory hormone hepcidin (1621 ± 329 pg/ml vs. 448 ± 171 pg/ml; p < 0.05), the iron-binding antimicrobial protein lipocalin-2 (LCN2; 10,494 ± 474 pg/ml vs. 4667 ± 166 pg/ml; p < 0.05), and the ferroxidase ceruloplasmin (429.2 ± 21.9 ng/ml vs. 358.4 ± 11.1 ng/ml; p < 0.05) were all significantly elevated in IAV-infected mice relative to naïve controls (Figure 4C–E). The coordinated induction of these iron-sequestration mediators suggests that the host mounts an active nutritional immunity response early during IAV infection – likely as a protective mechanism to limit microbial access to this essential metal. Notably, although IAV elicited robust induction of these iron-responsive mediators relative to naïve mice, their expression was only modestly altered by secondary *Af* challenge (Supplemental Figure 4), reinforcing the interpretation that IAV is the primary driver of pulmonary iron-associated inflammation and establishing that elevated iron availability is present in the lung at the precis moment of *Af* conidial deposition.

### Elevated iron promotes Aspergillus fumigatus germination and impairs macrophage-mediated fungal killing

Having established that IAV infection generates a pulmonary iron-overloaded environment, we next investigated whether elevated extracellular iron directly promote *Af* pathogenesis – either by enhancing fungal growth or by impairing host immune cell function.

We first assessed whether elevated iron affected *Af* radial growth on solid medium supplemented with iron concentrations corresponding to those measured in naïve and IAV-infected BALF. No significant differences in colony diameter were observed over a 3-day period (Supplemental Figure 5), indicating that iron does not broadly enhance fungal proliferation under these conditions. We therefore shifted our focus to germination kinetics, as *in vivo* IAPA coinfection is associated with increased fungal germination [22], and germination represents a critical transition from immune-resistant conidia to invasive hyphae.

Using our established *in vitro Af* germination assay [47], we evaluated *Af* germination in liquid GMM medium containing iron concentrations matching either naïve or IAV-infected BALF. Under both conditions, germination commenced at approximately 5 hours and plateaued near 90% by 8 hours (Supplemental Figure 7). Strikingly, iron concentrations equivalent to those in IAV-infected airways significantly accelerated germination, with the greatest differential observed at 6 hours -- a 10% increase relative to naïve iron concentrations (38.7 ± 3.7% vs. 26.5 ± 3.6%; p < 0.05) (Figure 5A). This effect was recapitulated in more physiologically relevant lung homogenate (LH) medium, where differences between iron conditions were statistically significant as early as 4 hours post-inoculation (13.2 ± 1.0% vs. 8.4 ± 1.2%; p < 0.01) (Figure 5B).

**Figure 5.**
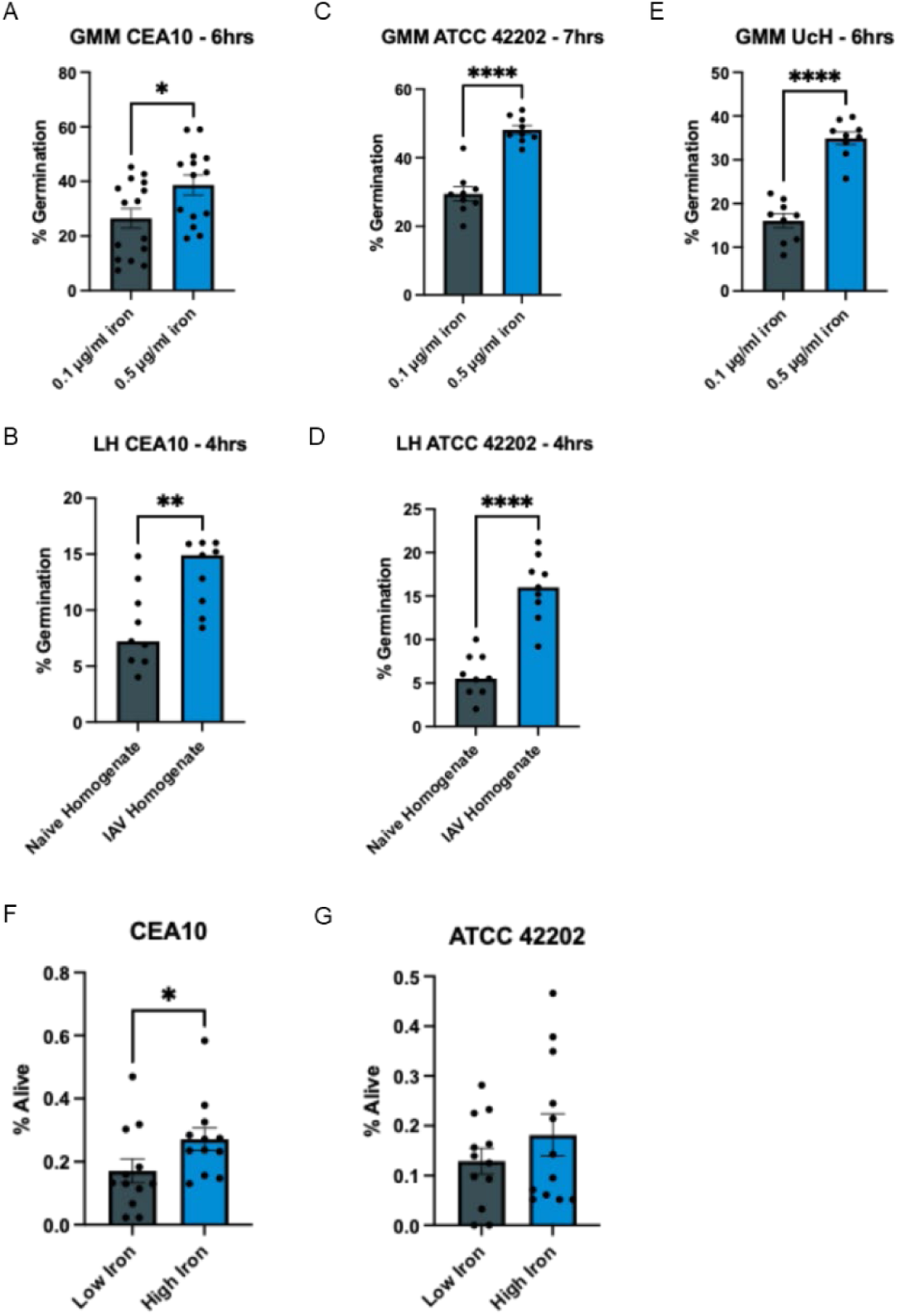
Iron overload promotes *Aspergillus fumigatus* germination and impairs macrophage-mediated fungal killing. *In vitro* germination of *A. fumigatus* strains (A) CEA10, (C) ATCC 42202, and (E) UCH cultured in glucose minimal medium (GMM) supplemented with high (0.5 mg/ml) or low (0.1 mg/ml) iron. Percent germination was quantified at 6 hours (CEA10 and UCH) or 7 hours (ATCC 42202) at the timepoint corresponding to the greatest difference in germination for each strain. Data in panel A are pooled from five independent experiments, while data in panels C and E are pooled from three independent experiments. *In vitro* germination of *A. fumigatus* strains (B) CEA10 and (D) ATCC 42202 in lung homogenate (LH) media. Percent germination was assessed at 4 hours post-inoculation. Data shown are pooled from three independent experiments. Macrophage-mediated killing of *A. fumigatus* (F) CEA10 and (G) ATCC 42202 using fetal-derived lung alveolar macrophages (FLAMs) under high- and low-iron conditions. Percent fungal survival was quantified by CFU enumeration following macrophage co-culture. Data are pooled from four independent experiments. Bars represent mean ± SEM, with individual points denoting biological replicates. Statistical significance was determined using unpaired two-tailed Mann–Whitney U tests. *p* < 0.05 (\**), p < 0.01 (**), p < 0.0001 (*\*\*\*\*)

To determine whether iron-enhanced germination was strain-specific or represented a general biological phenomenon, we extended these experiments to two additional *Af* strains: the ATCC 42202 laboratory strain which others have used in their model of IAPA coinfection [48–50] and a novel clinical isolate (UCH) obtained directly from an IAPA patient. Notably, in our hands both the ATCC 42202 and UCH *Af* isolates elicited cytokine responses comparable to those induced by CEA10 in the murine IAPA model (Supplemental Figure 7A-E), demonstrating similar regulation of inflammatory mediators across genetically distinct *Af* isolates. Most importantly, significant fungal germination of both the ATCC 42202 and UCH *Af* isolates was observed in the murine IAPA model, similar to what was observed with CEA10 (Supplemental Figure 7F). In GMM medium, ATCC 42202 showed significantly increased germination under high-iron concentrations across all timepoints, with a peak ∼20% increase at 7 hours (48.1 ± 1.2% vs. 29.4 ± 2.1%; p < 0.0001) (Supplemental Figure 6 and Figure 5C). UCH germination was similarly enhanced at 6 hours (34.9 ± 1.4% vs. 16.0 ± 1.6%; p < 0001) (Figure 5E and Supplemental Figure 7). Consistent results were observed in LH media, where ATCC 42202 exhibited significantly elevated germination under high-iron conditions compared to naïve iron concentration with approximately 10% increased germination at 4 hours (15.9 ± 1.2% vs. 5.9 ± 0.8%; p<0.0001) (Figure 5D). The reproducibility of these effects across three genetically diverse *Af* strains – including a patient-derived clinical isolate – strongly supports the conclusion that iron-enhanced germination is a conserved, strain-independent phenotype.

We next examined whether elevated iron concentrations also impaired host macrophage antifungal function. For these experiments, macrophages were pre-incubated overnight with iron dextran at concentrations corresponding to naïve (0.1 mg/ml) or IAV-infected (0.5 mg/ml) lung conditions – concentrations selected to maintain the approximately 5-10-fold differential observed in BALF while reflecting the tissue iron levels encountered by myeloid cells in the context of hemorrhagic lung injury and hemoglobin-associated iron sources [51–54]. Macrophages were subsequently challenged with CEA10 or ATCC 42202 resting conidia -- to best model the initial interactions occurring *in vivo –* at an MOI of 1, and fungal survival was assessed by CFU enumeration following an 8-hour co-culture.

Across both *Af* strains, significantly greater CFUs were recovered from macrophages pre-incubated under high-iron concentrations compared to lwo-iron concentration (Figure 5F-G), confirming that iron overload directly and substantially impairs macrophage antifungal killing capacity. These *in vitro* findings closely recapitulate the phagocytic function observed in IAPA coinfected mice *in vivo* [22], and are consistent with prior mechanistic studies demonstrating that elevated iron uncouples phagocytosis from fungicidal activity – associated with decreased lysosomal acidification, compromised membrane integrity, and reduced macrophage viability [53].

Collectively, these data demonstrate that the iron-enriched pulmonary environment created by IAV infection simultaneously accelerates *Af* germination – facilitating the transition to invasive hyphal growth – and directly suppresses macrophage-mediated fungal killing, providing a compelling dual mechanism for increased IAPA susceptibility in the context of prior IAV infection. These findings are also consistent with observations in severe SARS-CoV-2 infection, where excess airway iron has similarly been shown to enhance fungal growth while impairing neutrophil antifungal activity [55], suggesting that elevated airway iron may represent a shared, infection-agnostic mechanism predisposing to secondary *Af* infection across viral pneumonias.

## Discussion

In this study, we combined single-cell transcriptomics, functional *in vitro* assays, and cross-species comparative analysis to comprehensively define the immunological and metabolic landscape of IAPA in a validated murine model. Our findings reveal a coordinated disruption of pulmonary monocyte and macrophage function during IAPA coinfection, mechanistically driven in part by iron overload, impaired interferon signaling, and reduced responsiveness to inflammatory cues. These data provide important insight into why fungal clearance fails following prior viral infection and identify conserved transcriptional defects that parallel those described in human IAPA patients [30, 31].

A central finding of this study is that our murine IAPA model is associated with a fundamental remodeling of the myeloid transcriptional landscape -- a shift away from interferon-driven antiviral immunity toward tissue injury, stress adaptation, and metal-handling signatures. The transcriptional reprogramming observed in this study closely mirrors observations in human IAPA, where reduced type I and type II IFN responsiveness (including attenuated expression of *STAT1, STAT2, ISG15, MX1, IFNGR1,* and *CEBPB*) together with impaired neutrophil effector functions (including impaired *CTSA, CTSD, CTSS*, *S100A4, S100A10,* and *S100A11*) characterize the IAPA immune environment [30]. Reactome and ClueGO pathway analyses in our murine IAPA model consistently identified suppression of type I and type II interferon responses, degranulation, and viral defense programs in the IAPA coinfection cohort, providing strong cross-species validation of these transcriptional defects.

Module scoring further revealed that IAPA coinfected myeloid cells exhibit impaired activation programs encompassing ROS production, phagocytosis, IL-1β signaling, and TNF-α responsiveness. Strikingly, myeloid cells from IAPA coinfected mice also displayed transcriptional evidence of a metabolic shift toward oxidative phosphorylation at the expense of glycolysis -- a pattern reported in human CAPA macrophages and associated with defective *Aspergillus* spp. killing [31]. This metabolic reprogramming, which moves cells away from the mTOR-dependent glycolytic program required for efficient antifungal defense, represents a potentially targetable metabolic vulnerability in IAPA.

We demonstrated that IAV infection produces significantly elevated total iron levels in the airways by day 6 post-infection -- the precise time point at which secondary *Af* infection occurs in our model. This iron overload was accompanied by coordinated induction of iron-responsive mediators, including IL-6, hepcidin, LCN2, and ceruloplasmin, consistent with the activation of a host nutritional immunity response. However, a critical and paradoxical finding was the disconnect between extracellular iron burden and cellular iron-handling capacity during IAPA coinfection: despite the iron-enriched pulmonary environment, myeloid cells in IAPA coinfected lungs showed marked downregulation of iron uptake, import, storage, and sequestration pathways -- including reduced expression of *Fth1*, *Ltf, and Lcn2*. This failure of iron-regulatory adaptation during coinfection likely reflects the broader immune exhaustion and transcriptional reprogramming that characterizes IAPA and represents a permissive condition that simultaneously enriches the extracellular iron available for fungal exploitation while leaving the-mediated damage response unchecked.

By functionally characterizing the consequences of this iron overload, we found that elevated iron concentrations equivalent to those observed in IAV-infected airways both accelerated *Af* gemmation across three genetically diverse strains and significantly impaired macrophage-mediated fungal killing. The *in vitro* macrophage killing findings recapitulate the impaired phagocytic function seen in IAPA coinfected mice [22] and suggest that iron directly suppresses the antifungal effector capacity of pulmonary myeloid cells. Elevated iron has been associated with increased susceptibility to invasive mold infections in diverse clinical contexts, including patients with hepatic iron overload and lung transplant recipients [36, 44], and experimental models have demonstrated that increased tissue iron promotes *Af* invasion via enhanced allograft colonization and augmented invasiveness following local iron supplementation [54]. Prior work also supports the role of iron overload in impairing macrophage antifungal functions, while increased iron concentrations enhanced conidial uptake, they impaired intracellular fungal killing, which was associated with decreased lysosomal acidification, membrane instability, and reduced macrophage viability [53]. Our findings extend this body of evidence to the specific context of viral coinfection and, together with parallel observations in SARS-CoV-2 infection [55], suggest that excessive airway iron may represent a conserved, virus-agnostic mechanism of secondary fungal susceptibility.

Our findings have direct translational implications. The reduced IL-1β signaling observed in our murine IAPA model mirrors the functional deficits reported in human BALF [30], providing a mechanistic rationale for therapeutic restoration of the IL-1R axis. Indeed, the IL-1R antagonist Anakinra has demonstrated clinical promise by rebalancing the inflammatory environment and promoting adequate ROS-mediated antifungal defense [56]. Aerosolized Toll-like receptor (TLR) agonists such as Pam2-CSK4 have shown efficacy in priming airway epithelial immune recognition to enhance antifungal defense in IAPA models [57]. Additionally, targeted early dampening of excessive IFN-γ signaling via Emapalumab represents a promising strategy to preserve alveolar macrophage function, prevent successive immunosuppressive events, and restore ROS-dependent *Af* killing [23]. Importantly, our identification of iron dysregulation as a key disease-initiating factor in IAPA opens new therapeutic possibilities: interventions that restore iron sequestration — such as iron chelation therapy — could represent a novel host-directed strategy to limit the risk of secondary aspergillosis in patients with severe respiratory virus infections. Future studies should rigorously evaluate whether iron chelators can be safely deployed in this context and whether iron restriction is sufficient to restore antifungal myeloid cell function.

It is also relevant to note that SARS-CoV2-infected patients treated with tocilizumab – an IL-6 receptor antagonist -- may face elevated risk of developing CAPA [11]. Given the central role of IL-6 in regulating both inflammatory responses and iron homeostasis through hepcidin induction, understanding the consequences of IL-6 pathway inhibition for pulmonary iron regulation and fungal susceptibility will be important in informing treatment decisions in severe respiratory virus infection [11].

Several limitations of the present study merit acknowledgement. First, direct comparisons between murine and human single-cell transcriptomic datasets require caution given species-specific differences in immune cell subsets, gene regulation, and lung architecture. Second, our analysis was restricted to the interstitial macrophages and monocytes and did not include alveolar macrophages, neutrophils, or lymphoid populations that may independently contribute to IAPA pathogenesis. Third, while we identified key transcriptional and functional defects relating to iron metabolism and phagocyte dysfunction, we did not directly interrogate the molecular mechanisms linking iron overload to impaired interferon signaling and immune exhaustion – are important area for future mechanistic investigation. Fourth, although we demonstrate that elevated airway iron is sufficient to promote *Af* germination and impair macrophage killing, our studies do not formally demonstrate that this iron overload is necessary for IAPA susceptibility; future experiments employing iron chelation *in vivo* will be required to address this causal question. Finally, our analysis focused on early infection time points that capture the critical window of secondary *Af* susceptibility but does not address immune dynamics during disease progression or resolution. Longitudinal studies encompassing later time points will be needed to fully characterize the arc of IAPA coinfection.

In summary, our study identifies a coordinated immunometabolic vulnerability in IAPA coinfection in which prior IAV infection primes the lung for fungal overgrowth through four interconnected mechanisms: (1) induction of local iron overload; (2) paradoxical impairment of myeloid iron-handling capacity; (3) suppressing cytokine and interferon signaling; and (4) metabolic reprogramming of immune cells toward a less responsive, oxidative phosphorylation-dependent state. These combined changes create a pulmonary environment that simultaneously promotes *Af* germination and shields the fungus from effective immune clearance. Given the increasing incidence and persistently high mortality of IAPA, particularly in ICU settings, our findings underscore the urgent need mechanistically informed interventions – including strategies that target iron dysregulation, restore immunometabolic fitness, and rescue interferon responsiveness in the vulnerable post-viral lung.

## Materials and Methods

### *Aspergillus fumigatus* strains and culture conditions

*A. fumigatus* strains CEA10, ATCC 42202 and UCH were used throughout this study. The UCH clinical isolate was obtained from the UC Health Clinical Microbiology Laboratory at the University of Cincinnati Medical Center. The deidentified isolate was released to Dartmouth College under a material transfer agreement. Strains were maintained on 1% glucose minimal medium (GMM) agar plates. For conidia collection, plates were incubated for 3 days at 37°C. Conidia were harvested by gently scraping plates into PBS containing 0.01% Tween-80, filtered through sterile Miracloth, washed, and resuspended in PBS. Conidia were enumerated using a hemocytometer and stored in PBS at 4°C until use.

### Murine model of influenza-associated pulmonary aspergillosis (IAPA)

Female C57BL/6J mice (Jackson Laboratory), 10–15 weeks of age, were used for all *in vivo* studies. To model IAPA, mice were anesthetized with isoflurane and intranasally inoculated with 100 EID50 of the A/Puerto Rico/8/34 strain of IAV (Charles River Laboratories). Six days after IAV challenge, mice were anesthetized with isoflurane and intratracheally challenged with 100 µL PBS containing 3.5 × 10^7^ *A. fumigatus* CEA10 conidia. Four groups of mice were studied, which included PBS/PBS (naïve), IAV/PBS (IAV), PBS/*Af* (CEA10), and IAV/*Af* (IAPA coinfection) conditions as indicated. Mice were euthanized 24 h after fungal challenge (day 7 post-IAV challenge) for downstream analyses unless otherwise specified. All procedures were approved by the Dartmouth College Institutional Animal Care and Use Committee (IACUC, Protocol #00002243).

### Bronchoalveolar lavage fluid (BALF) collection and soluble mediator quantification

BALF was collected 24 h after fungal challenge by lavaging lungs with 2 ml PBS containing 0.05*M* EDTA. BALF was clarified by centrifugation, and supernatants were stored at −20°C until analysis. IL-6 was quantified by ELISA (BioLegend) and iron-regulatory proteins hepcidin, ceruloplasmin, and lipocalin-2 (LCN2) were quantified by ELISA (R&D Systems) according to manufacturers’ instructions.

### Bronchoalveolar lavage fluid (BALF) collection and trace metal quantification

BALF was collected 6 d after IAV challenge by lavaging lungs with 2 ml PBS. Cell-free BALF supernatant is then collected following centrifugation (5 min, 1500 rpm, 4°C) and supernatants were stored at −20°C until analysis. Trace metal levels within the cell-free BALF supernatant were analyzed by ICP-QQQ-MS (8900 Triple Quadrupole ICP-MS, Agilent, Wilmington, DE) using a prepFAST M5 autosampler (ESI, Omaha, NE) by the Dartmouth Trace Element Analysis Core. Iron, cooper and zinc were analyzed in helium gas mode to reduce polyatomic interferences. Both ^56^Fe and ^57^Fe were measured during the analysis and the concentrations for these two isotopes were in good agreement being 2.6% average relative difference which is within the precision of daily instrument performance and strongly suggests no effects of polyatomic interferences on Fe measurement. A secondary source standard was used as an initial calibration check and every ten samples and average recovery for Fe was 97 ± 4%.

### Lung cell isolation and fluorescence-activated cell sorting (FACS)

At 24 h post-fungal challenge (day 7), mice were euthanized, lungs were perfused, excised, minced, and digested in RPMI 1640 containing 1 mg/ml collagenase D (Roche) for 30 min at 37°C with agitation. Digested tissue was passed through a 70 µm mesh filter, subjected to red blood cell lysis, and resuspended in FACS buffer (PBS containing 2% FBS and 2 mM EDTA). Cells were enriched for CD11b^+^ cells by magnetic separation (Miltenyi Biotec) prior to sorting.

For sorting monocytes and interstitial macrophage populations used for scRNA-seq, live immune cells were isolated using a multi-step strategy. Cells were first CD45-enriched, followed by exclusion of intravascular CD45^+^ cells (BD Biosciences; A20) (intravenous CD45 labeling), and depletion of lineage markers including Ly6G (Biolegend; 1A8), NK1.1 (Biolegend; PK136), CD4 (Biolegend; GK1.5), CD8a (Biolegend; 53-6.7), Siglec-F (Biolegend; S17007L), and CD19 (Biolegend; 6D5) to remove neutrophils, NK cells, T cells, alveolar macrophages, and B cells. Monocytes/interstitial macrophages were then positively selected as CD45^+^ (Biolegend: 30-F11) CD64^+^ (Biolengend: X54-5/7.1) CD11b^+^ (Invitrogen: M1/70) cells. A live/dead viability dye (DAPI: D9542) was included immediately prior to sorting to ensure collection of viable cells only. Sorting was performed at 4°C using a BD FACSAria II (BD Biosciences).

### Single-cell RNA sequencing and computational analysis

Sorted CD64^+^ CD11b^+^ monocytes and interstitial macrophages were processed using the Chromium Next GEN Single Cell 3′ v3.1 platform (10x Genomics) following the manufacturer’s protocol. Libraries were sequenced on an Illumina NovaSeq 6000 at a target depth of ∼30,000 reads per cell. Raw base call files were processed using Cell Ranger v7.0 against the mm10 reference genome.

Downstream analysis was performed in R (v4.4.1) using Seurat (v5.3). Cells were filtered to remove low-quality droplets and potential doublets, excluding cells with < 200 detected genes or > 10% mitochondrial transcripts. Data were normalized using SCTransform and integrated across conditions using canonical correlation analysis (CCA). Clusters were identified by graph-based clustering. Cell–cell communication inference was conducted using CellChat, with analyses performed per condition and compared across conditions.

### Differential expression and pathway enrichment analyses

Differential expression between conditions was performed using Seurat’s FindMarkers function (Wilcoxon rank-sum test) using all expressed genes were retained (*min.pct* = 0, *logfc.threshold* = 0) to enable unbiased downstream pathway analyses. P-values were adjusted for multiple hypothesis testing using the Benjamini–Hochberg false discovery rate (FDR) correction, and genes with adjusted *p* < 0.05 were considered significantly differentially expressed.

Gene Ontology (GO) and pathway enrichment analyses were performed using clusterProfiler (v4.6) and fgsea with enrichment significance determined using FDR-adjusted p-values. Gene set enrichment analysis (GSEA) was performed using Reactome pathway annotations, leveraging ranked gene lists derived from differential expression analyses. We accessed the enrichment scores for pathways with minimum 15 and maximum 500 genes shared with our data, and minimum 15% pathway coverage. Pathway-level analyses were additionally performed on all negative differentially expressed genes (DEGs) from IAPA vs. IAV cells using ClueGO (v2.5.9) within Cytoscape. ClueGO analyses incorporated both GO Biological Process and Reactome databases, using a right-sided hypergeometric test with Benjamini–Hochberg correction. Semantic redundancy among GO terms was reduced using rrvgo based on pairwise term similarity. Pathways with minimum 3 genes or 2% of pathway genes overlap with the input DEG were selected. Selective pathways analyses were performed by filtering database terms to reflect pathways of interest. GO terms were filtered for iron related pathways and Reactome pathway results were filtered to include pathways reported in human BALF datasets.

### Module Scoring

Curated gene sets representing specific biological processes (e.g., iron storage, iron uptake, iron sequestration, glycolysis, oxidative phosphorylation, mTOR signaling, phagocytosis, reactive oxygen species production, macro/micronutrient metabolism and cytokine-responsive programs) were assembled based on established literature and pathway databases (see appendix). These gene sets were used to calculate single-cell module scores using Seurat’s AddModuleScore function. Module scores were compared across conditions using violin plots. Means and standard error of mean (SEM) were calculated in R. Statistical comparisons of module scores between conditions were performed using non-parametric tests (Wilcoxon rank-sum)

### In vivo germination assays in defined media and lung homogenate (LH)

For *in vitro* germination assays, conidia were inoculated at 1 × 10^7^ conidia/mL into 2 ml of media in glass 20 ml scintillation vials (VWR). Germination was assessed in GMM diluted into iron-free GMM to achieve final iron conditions corresponding to high-iron (0.5 µM) and low-iron (0.15 µM). For lung homogenate (LH) germination assays, LH media was prepared from either day 6 IAV-infected or naïve lungs, as previously described [47]. Cultures were incubated at 37°C with shaking at 300 rpm. Beginning at 3 h post-inoculation, wet mounts were prepared hourly and samples were vortexed with 1.0 mm beads to disrupt clumps prior to mounting. Germination was quantified manually using a 40× objective by counting ≥ 100 total conidia and germlings per sample at each time point.

### Macrophage antifungal killing assays under iron modulation

To assess antifungal effector function under defined iron conditions, fetal liver–derived alveolar macrophages were differentiated in the presence of GM-CSF (30 ng/mL) and TGFβ (15 ng/mL) under standard culture conditions, as previously described [58]. Macrophages were seeded in 24-well plates and allowed to adhere overnight at 37°C with 5% CO2. Cells were pretreated for 16 h with iron dextran (D8517; Millipore Sigma) at 0.15 µM (low iron) or 0.5 µM (high iron) in complete media. Conidia (CEA10 or ATCC 42202) were added at 1 × 10^6^ conidia/mL (MOI 1) and incubated for 1 h at 37°C for uptake. Wells were washed with PBS to remove non-adherent conidia, and intracellular killing was assessed after 1 h and 8 h. Cells were lysed with 10% SDS, and released fungal material was serially diluted and plated onto GMM agar. After 48 h incubation at 37°C, CFUs were enumerated and expressed as percent survival relative to input where indicated.

### Statistical analysis

Statistical analyses were performed in Prism v10/R 4.4.1. For single-cell RNA-sequencing analyses, statistical testing was conducted within the Seurat framework unless otherwise specified. DEG analyses were performed using the Wilcoxon rank-sum test with both DEG and pathway analyses using the Benjamini–Hochberg false discovery rate (FDR) correction. Module scores were compared using Wilcox rank-sum or Kruskal–Wallis tests if more than two conditions were graphed. Cell–cell communication differences inferred by CellChat were assessed using permutation-based statistical testing implemented within the CellChat framework. Differences in interaction number and interaction strength across conditions were evaluated using the default bootstrapping procedures provided by the package.

Data is presented as mean ± SEM unless otherwise stated. The number of biological replicates and independent experiments is indicated in the corresponding figure legends. comparisons between groups were performed by Mann–Whitney U tests were used. Statistical significance was defined as follows: p < 0.05 (*), p < 0.01 (**), p < 0.001 (***), and p < 0.0001 (****).

### Data availability

The sequencing raw data and processed data used in this article are available in the Gene Expression Omnibus under accession code GSE328605.

## Acknowledgements

We thank Gary Ward for flow cytometry support, Owen Wilkins for sequence processing support, and Claudia Jakubzick for consultation on myeloid cell sorting. This work was supported in part by National Institutes of Health (NIH) grant R21 AI152019 (JJO). MSG was supported by T32 AI007363 (PI: Claudia Jakubzick). NIH grants P20-GM130454, S10-OD025235, and S10-OD030242, which supports the single-cell genomics core, and NIH grant P30-CA023108, which supports the Dartmouth Cancer Center’s flow cytometry (RRID: SCR_019165), genomics (RRID: SCR_021293), and Trace Element (RRID: SCR_009777) shared resources.

