## Supplement Figures for "Myeloid Cell States in Influenza-Associated Pulmonary Aspergillosis Are Shaped by Iron Overload and Metabolic Reprogramming"

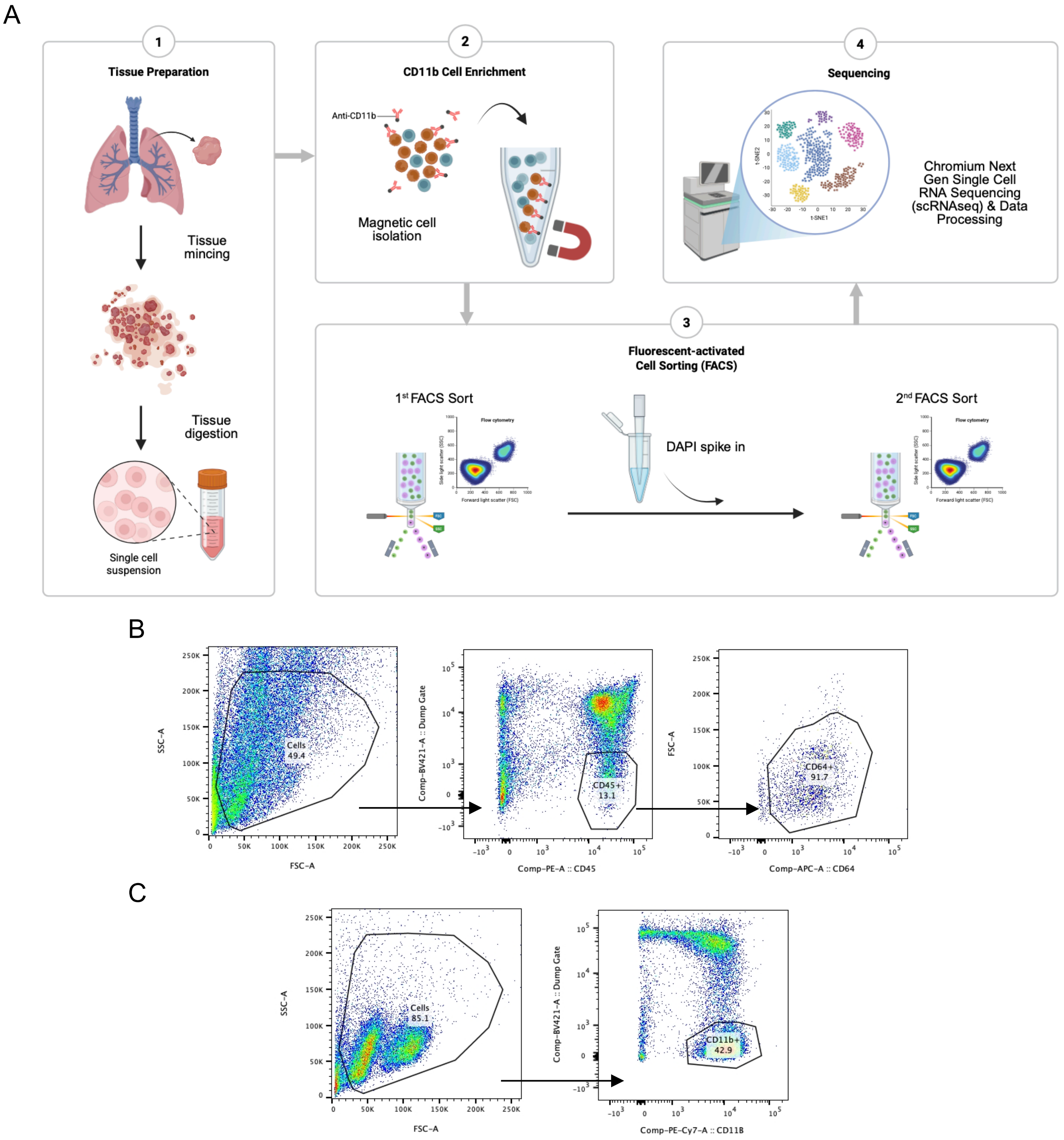


**Supplemental Figure 1. Two-step FACS sorting workflow used for isolation of lung monocyte/macrophage populations for scRNA-seq.**

(A) Schematic overview of the cell isolation workflow. Lung tissue was mechanically dissociated and digested to generate a single-cell suspension, followed by CD11b-positive magnetic enrichment. Enriched cells then underwent a two-step fluorescence-activated cell sorting (FACS) strategy prior to Chromium Next Gem single-cell RNA sequencing and downstream data processing. DAPI was added between the first and second sort as a viability dye. (B) Representative flow cytometry plots from the first-round sort. Total cells were initially gated by forward and side scatter, followed by CD45+ leukocyte selection and CD64+ enrichment. (C) Representative flow cytometry plots from the second-round sort performed on the collected population after DAPI addition. Cells were re-gated by forward and side scatter, and DAPI fluorescence, detected in the dump channel, was used to exclude nonviable DAPI-positive events during resorting. The final live CD11b+ population was collected for 10x Genomics single-cell RNA sequencing.

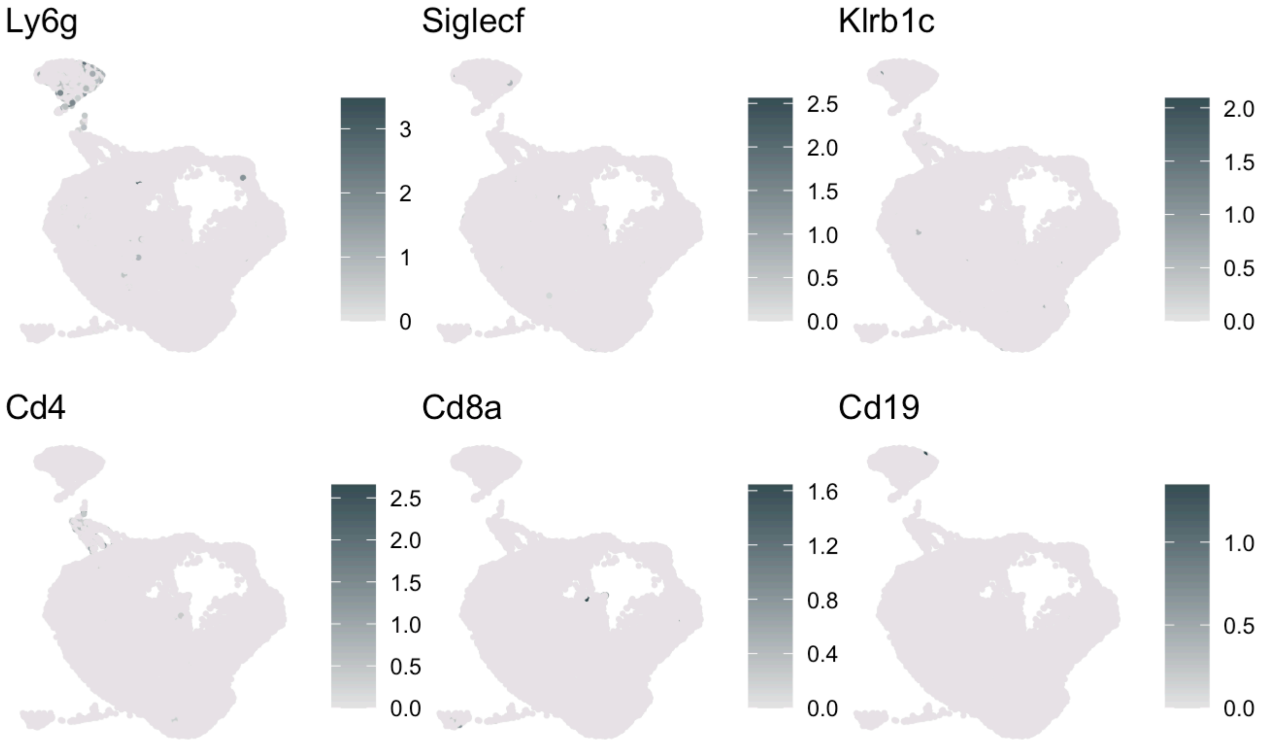


**Supplemental Figure 2. Feature plots of excluded lineage markers across the integrated sorted-cell dataset confirms successful enrichment prior to single-cell RNA sequencing.** Feature plots projected onto the integrated UMAP of all combined samples show expression of genes corresponding to cell populations excluded during fluorescence-activated cell sorting prior to 10x Genomics single-cell RNA sequencing. Plotted genes include markers associated with neutrophils, NK cells, alveolar macrophages, T cells and B cells lineages incorporated into the dump-channel exclusion strategy. The overall low or restricted expression of these genes across the dataset confirms effective depletion of unwanted cell populations during sorting and enrichment of the target monocyte/macrophage compartment before sequencing.

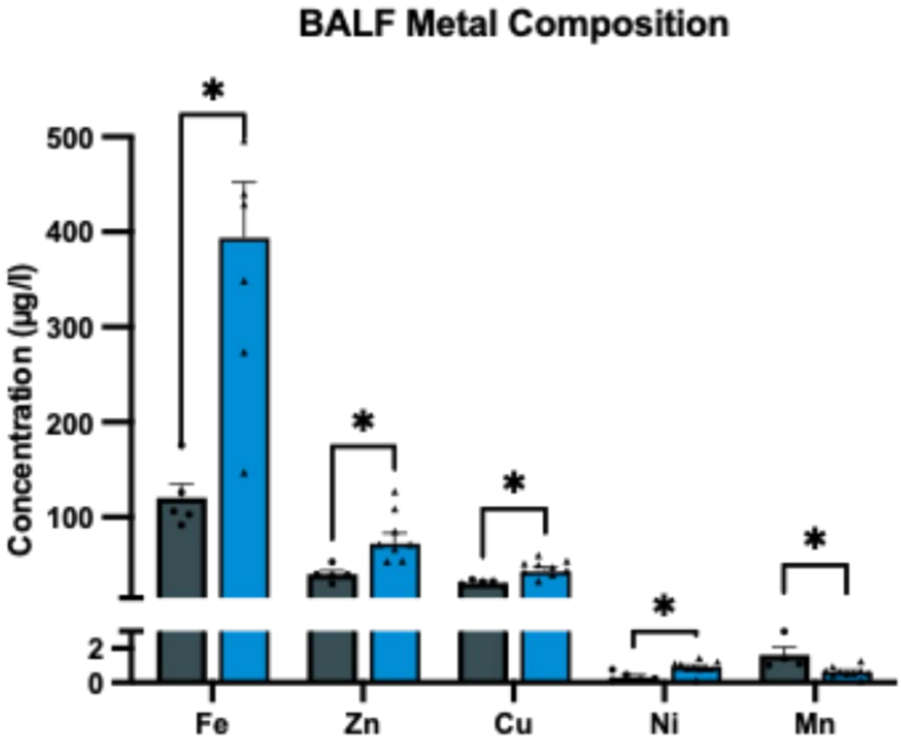


**Supplemental Figure 3. Metal composition of bronchoalveolar lavage fluid in naïve and IAV-infected mice.**
BALF concentrations of iron (Fe), zinc (Zn), copper (Cu), nickel (Ni), and manganese (Mn) measured in naïve and IAV-infected mice. Data are shown from a single representative experiment (n = 5 naïve, n = 10 IAV). Bars indicate mean ± SEM. Statistical comparisons for each metal were performed using a Mann-Whitney test.

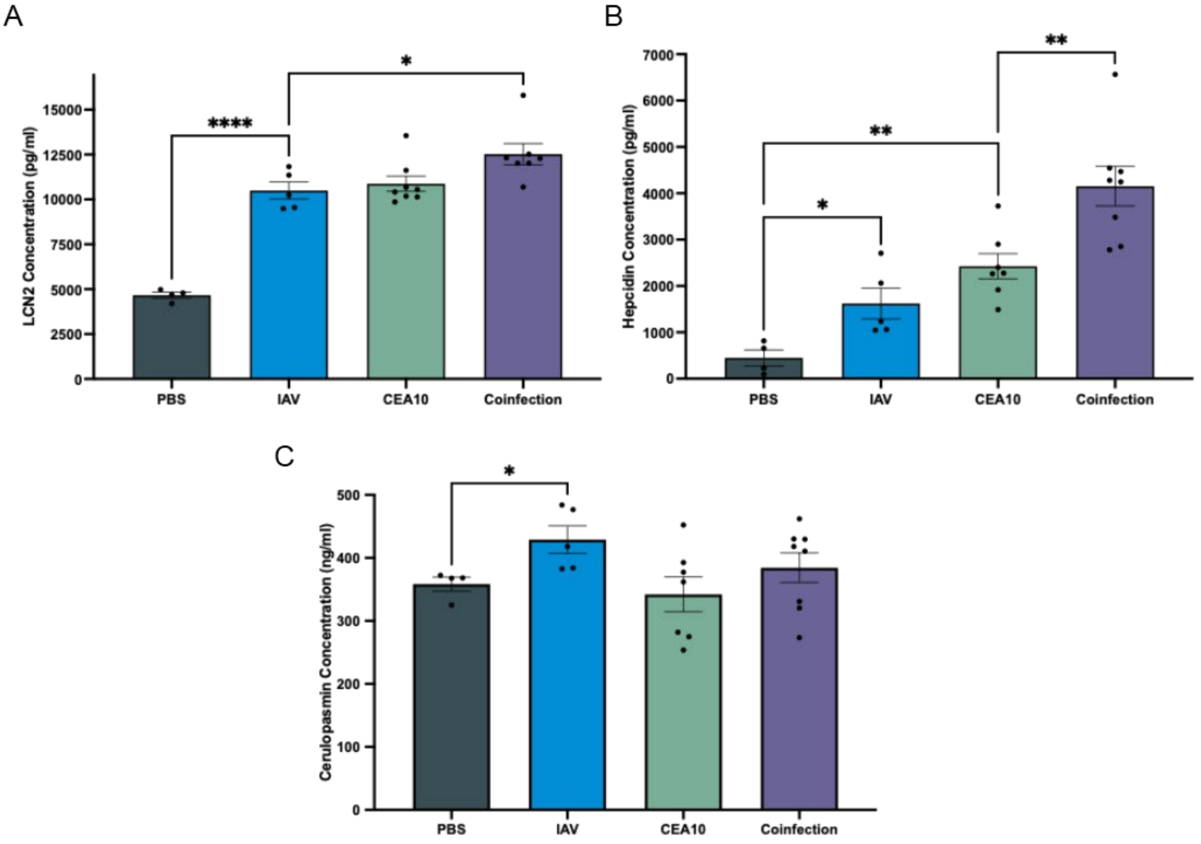


**Supplemental Figure 4. ELISA-based quantification of iron-regulatory proteins.**
(A) Lipocalin-2 (LCN2), (B) hepcidin, and (C) ceruloplasmin concentrations measured by ELISA in samples from four experimental conditions. Bars indicate mean ± SEM. Differences among groups were assessed by one-way ANOVA.

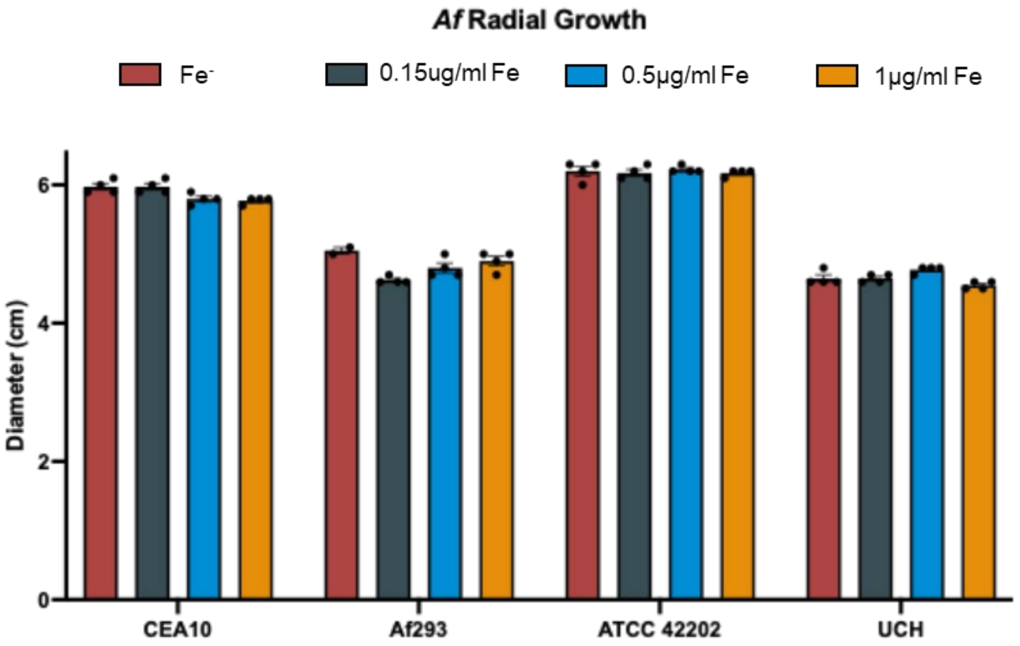


**Supplemental Figure 5. Radial growth of *Aspergillus fumigatus* strains across iron concentrations.**

Radial growth of CEA10, Af293, ATCC 42202, and UCH *A. fumigatus* strains on glucose minimal medium (GMM) agar formulated with increasing iron concentrations. Red bars represent growth on iron-free GMM. Grey and blue bars correspond to iron concentrations found in BAL of naïve (0.15 µg/mL) and IAV (0.5 µg/mL) mice, generated by mixing iron-free GMM with standard GMM. Orange bars correspond to standard GMM (1 µg/mL iron). Bars represent mean ± SEM.

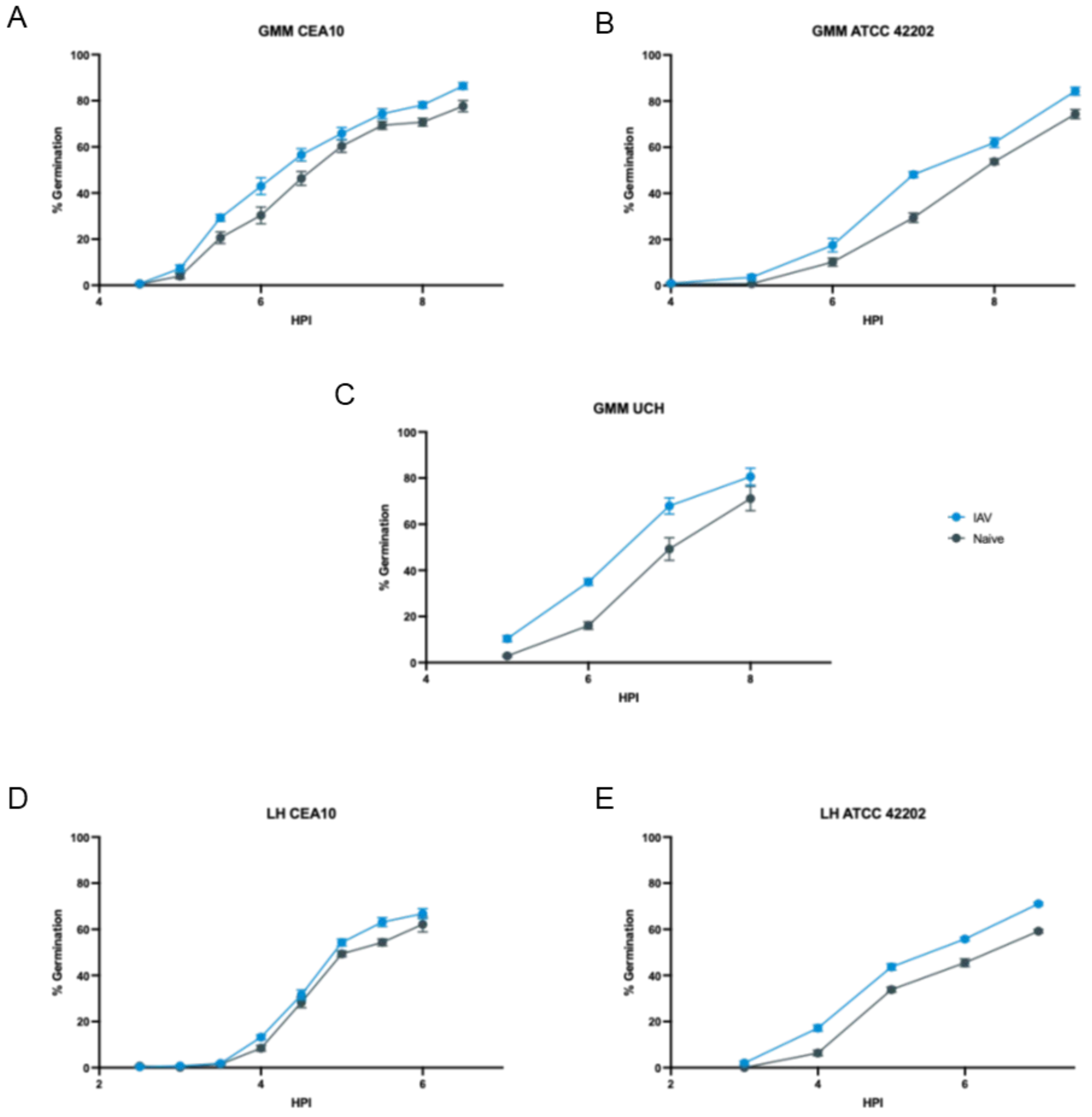


**Supplemental Figure 6. Germination curves of multiple *Aspergillus fumigatus* strains under iron-matched infection conditions.**

Germination kinetics of (A) CEA10, (B) ATCC 42202, and (C) UCH were measured in glucose minimal medium (GMM) adjusted to iron concentrations corresponding to BAL from naïve (0.15 µg/mL) and IAV-infected (0.5 µg/mL) mice by mixing iron-free GMM with standard GMM. Germination kinetics of (D) CEA10 and (E) ATCC 42202 were also measured in lung homogenate (LH) media derived from naïve and IAV-infected mice. Percent germination was quantified over time and plotted as hours post-inoculation (HPI). Data were pooled from multiple independent experiments, with each experiment containing three biological replicates. Points represent mean ± SEM.

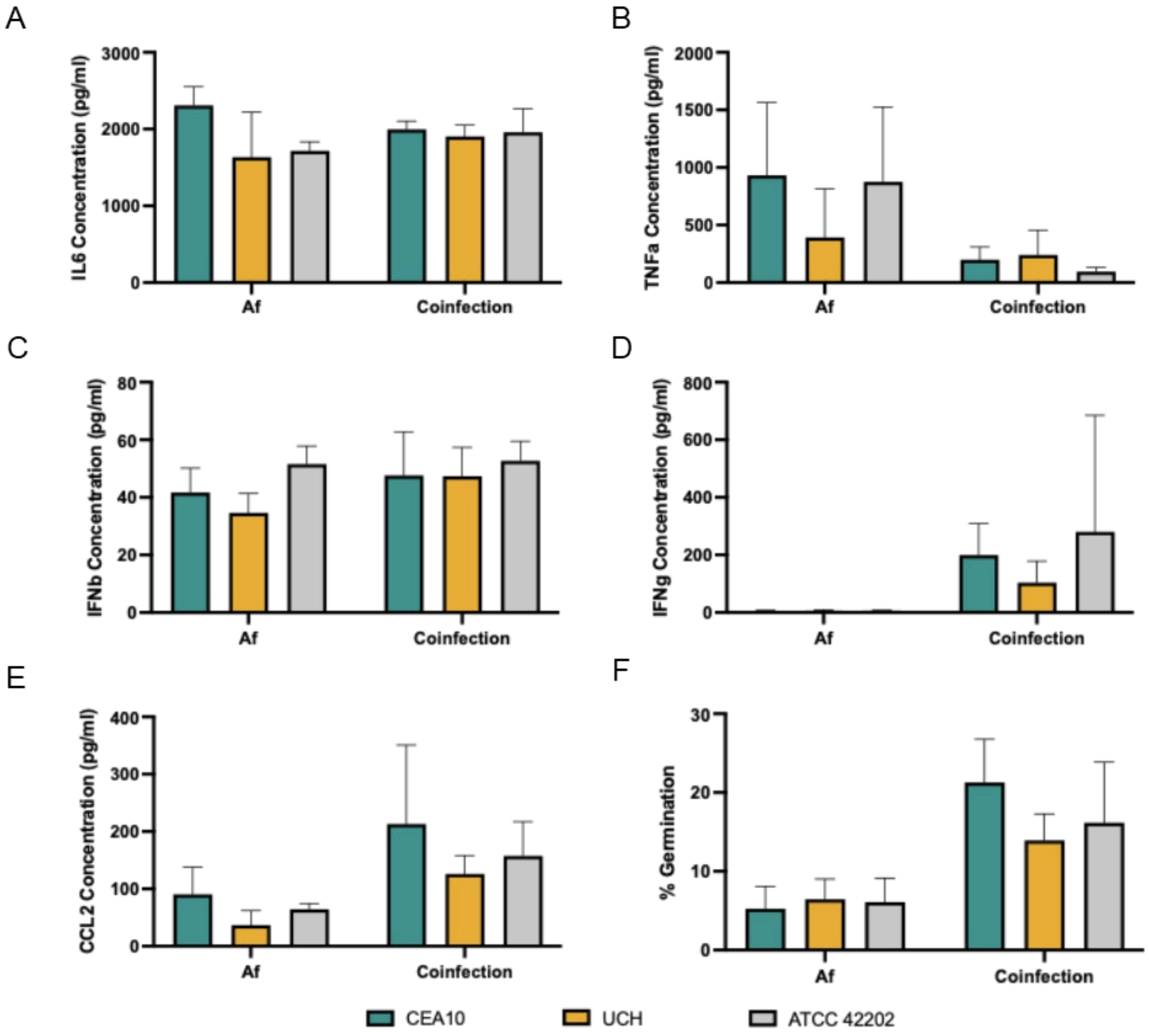


**Supplemental Figure 7. IAPA *A. fumigatus* strains exhibit comparable inflammatory and germination profiles in fungal infection alone and during coinfection.**

Cytokine production and fungal germination were compared among three *A. fumigatus* strains, CEA10, UCH, and ATCC 42202, during *Af* infection alone and coinfection conditions. (A) IL6, (B) TNFα, (C) IFNβ concentration, (D) IFNγ, and (E) CCL2 concentrations were determined by ELISA. (F) Percent germination was enumerated from GMS staining. Bars represent mean ± SEM.
